# Human gut archaea collection from Estonian population

**DOI:** 10.1101/2024.12.09.627479

**Authors:** Kateryna Pantiukh, Elin Org

**Author notes:** Corresponding authors: Elin Org and Kateryna Pantiukh.

## Abstract

While microbiota plays a crucial role in maintaining overall health, archaea, a component of microbiota, remain relatively unexplored. Here, we present a newly assembled set of archaeal metagenome-assembled genomes (MAGs) from 1,887 fecal microbiome samples. These archaeal MAGs were recovered for the first time from the Estonian population, specifically from the Estonian Microbiome Deep (EstMB-deep) cohort. In total, we identified 273 archaeal MAGs, representing 21 species and 144 strains (“EstMB MAGdb Archaea-273” MAGs collection). Of these 21 species, 12 species belonged to the order *Methanobacteriales* and *Methanomassilicicoccales*, other 9 species from *Methanomassiliicoccales* were novel. Notably, 7 of the 9 new species belonged to the *UBA71* genus. Given that the latest version of the Unified Human Gastrointestinal Genome (UHGG v2.0.2) database includes 27 archaeal species, we expanded the known archaeal diversity at the species level by 30%.

## Background & Summary

The human gut microbiota plays a crucial role in maintaining overall health, yet the diversity of archaea within it remains relatively unexplored. The last release of Unified Human Gastrointestinal Genome (UHGG) collection (v2.0.2) contains 28 archaeal species^1^. The dominant species in the human gut archaeome belonged to orders *Methanobacteriales* (87.15%) and *Methanomassiliicoccales* (12.43%), with much lower representation from *Methanomicrobiales* (0.26%) and *Halobacteriales* (0.17%). The most prevalent archaea species detected in the human gut is methanogenic archaea *Methanobrevibacter smithii*, belonging to the order *Methanobacteriales*, which is found in over 95% of adults^2^. Methanogenic archaea are functionally significant due to their capacity to consume molecular hydrogen, facilitate the efficient breakdown of organic substances, and contribute to processes like energy metabolism and adipose tissue deposition^3, 4^. Additionally, there is some evidence of diseases associated with altered methane production archaeal composition^5^. As a result, much of the research has primarily focused on methanogenic archaea, particularly the *Methanobrevibacter smithii* species^6,7^. However, numerous less common archaea species have remained undiscovered. Hence, further research especially from new populations is imperative to uncover a wider array of archaeal species present in the human gut, as they could potentially impact human health.

Our goal was to explore the diversity of archaea in the Estonian population and expand the existing collection of human gut archaeal genomes by identifying new species present in the Estonian population and potentially other closely related populations (**Figure 1a**). We utilized the Estonian Metagenome-Assembled Genomes collection (EstMB MAGdb) as a source of MAGs. This collection features a substantial number of MAGs, assembled thanks to the availability of high-coverage short-read sequencing data. A comprehensive overview of the collection, including data availability for 2,257 species-representative MAGs and the methods used for their assembly, is provided in Pantiukh et al., 2024^8^. The species-representative MAGs were generated by clustering an initial dataset of 84,762 bacterial and archaeal MAGs. Notably, no archaeal MAGs were retained in the final representative clusters, as they, along with low-quality bacterial MAGs, failed to cluster and filtered out during the process. This outcome prompted us to revisit the initial MAG dataset and specifically examine the archaeal MAGs, including those that failed to cluster. Upon reanalysis, we identified a significant number of high-quality archaeal MAGs among the previously unclustered MAGs. Here, we present this newly curated archaeal MAG dataset.

**Figure 1.**
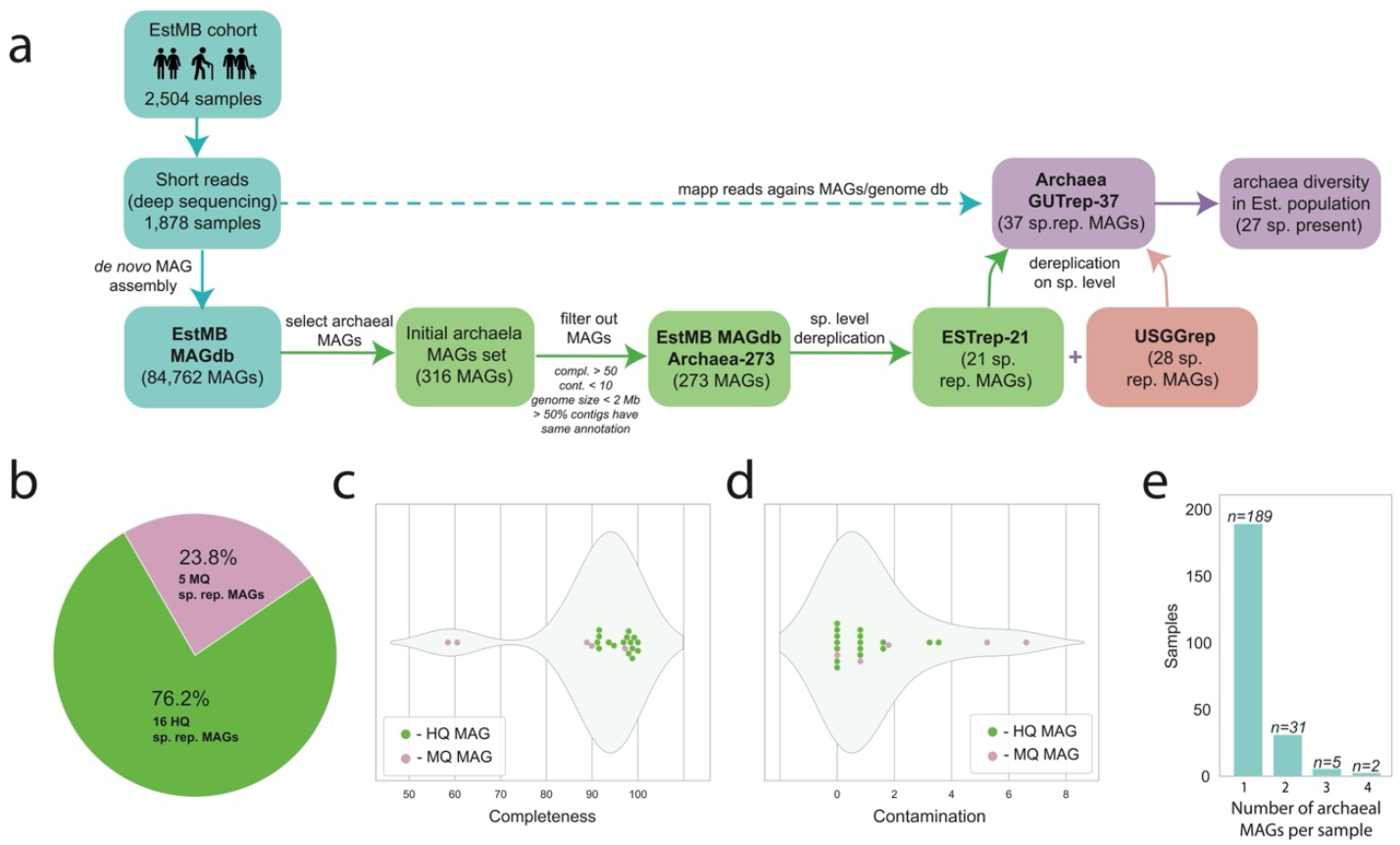
Overview of the creation of the Estonian archaeal MAGs collection, MAG quality assessment, and description of the origin samples. **a.** Study overview **b**. Quality estimation for the archaea MAGs from the “Archaea ESTrep-21” MAGs collection. High-quality MAGs (HQ)– 16; Medium-quality MAGs (MQ) – 5 **c**. Completeness of the MAGs from the “Archaea ESTrep-21” MAGs collection (%) **d**. Contamination of the MAGs from the “Archaea ESTrep-21” MAGs collection (%) **e**. Distribution of the number of MAGs per sample.

Samples from the Estonian population included individuals across various sex and age groups. Of these, 1,308 samples (69.64 %) were from females and 570 samples (30.35 %), from males. The participants had an average age of 50.05 years, with an age range from 23 to 89 years. The samples were sequenced with deep coverage, yielding an average of 16.49 ± 6.2 Gb of total base pairs per sample, or 56.13 ± 19.37 million paired reads per sample. Out of the l 84,762 recovered MAGs, we identified 316 archaeal MAGs using GTDB-tk **(Figure 1a)**. For all MAGs from this initial set, we evaluated completeness and contamination with CheckM2, genome size with seqkit and annotated each MAG contigs separately with GUNC. Low quality MAGs (completeness < 50%, contamination > 10%), MAG with abnormal genome size (>2Mb) and MAGs with non-constant contig annotation (<50% contigs have same archaeal annotation) were removed from final set, leaving 273 archaeal MAGs. We are referring to this final set of archaeal MAGs as “**EstMB MAGdb Archaea-273**” (**Supplementary Table S1**). For all MAGs from this set we performed gene prediction and annotation as well as tRNAs and clustered regularly interspaced short palindromic repeats (CRISPRS) prediction. The average number of coding sequences (CDS) per MAG is 1677 (1746 for *Methanomassiliicoccales* and 1540 for *Methanobacteriales*). Among these, on average 56% are unannotated hypothetical proteins (58% for *Methanomassiliicoccales* and 51% for *Methanobacteriales*). The average number of tRNA is 37. We were able to find from 0 to 3 CRISPRS per MAG.

Archaeal MAGs from “EstMB MAGdb Archaea-273” collection was clustered at the species level (ANI>95) and grouped into 21 species, with one representative genome selected from each species cluster. We refer to this final set of species representative archaeal MAGs as “**Archaea ESTrep-21**” MAG collection (**Supplementary Table S2**). Of the 21 species, 9 species (42.86%) could not be assigned to any previously described species. Interestingly, a large proportion of new species (7 out of 9 species) were affiliated with the genus *UBA71*, while the remaining 2 species belonged to the genera *Methanomassiliicoccus_A* and *DTU008*. All new species clusters contain 1 to 4 MAGs per cluster, except for the one new species cluster from the genus UBA71 (ESTrep_MAG_ID: R07_UBA71_undS_VLU9Q2), which has 11 MAGs in its species cluster. Known 12 species belong to two orders, most abundant in human gut: *Methanobacteriales* (83 MAGs, 3 known species) and *Methanomassiliicoccales* (190 MAGs, 9 known species). The Methanobacteriales order exclusively consists of species within the *Methanobrevibacter_A genus*, which includes *Methanobrevibacter_A woesei* (2 MAGs), *Methanobrevibacter_A smithii_A* (17 MAGs), and *Methanobrevibacter_A smithii* (64 MAGs). In contrast, the *Methanomassiliicoccales* order exhibits greater diversity and is represented by five distinct genera. The most prevalent genus, *UBA71*, contains four known species: *UBA71 sp905187815* (8 MAGs), *UBA71 sp016296195* (1 MAG), *UBA71 sp006954465* (29 MAGs), and *UBA71 sp006954425* (56 MAGs). The *Methanomethylophilus* and *MX-02* genera each include only a single known species: *Methanomethylophilus alvus* (41 MAGs) and *MX-02 sp006954405* (15 MAGs). The *Methanomassiliicoccus_A* genus comprises two known species: *Methanomassiliicoccus_A sp905203995* (3 MAGs) and *Methanomassiliicoccus_A intestinalis* (9 MAGs). The *DTU008* genus is represented by one known species, *DTU008 sp001421185*.

Out of 21 species representative MAGs, 16 (76.19%) had completeness greater than 90% and contamination less than 5% (HQ MAGs), while other 5 MAGs had completeness between 50% and 90% and contamination between 5% and 10% (MQ MAGs) (**Figure 1b**). These species representative MAGs from “**Archaea ESTrep-21**” MAG collection had a median completeness of 96.85% (Interquartile Range [IQR], 91.49 - 98.39) and a median contamination of 0.81% (IQR, 0.00 - 1.61) (**Figures 1c, Figure 1d**).

Typically, only one archaeal MAG is extracted from a sample. Among the 227 samples containing archaeal MAGs, 189 samples contain 1 archaeal MAG, 31 samples contain 2 archaeal MAGs, 5 samples contain 3 archaeal MAGs, and 2 samples contain 4 archaeal MAGs (**Figure 1e**).

Given the complexity of metagenome assembly, we anticipated that not all species present in the population would have corresponding MAGs. To gain a more comprehensive understanding of archaeal diversity in our population, we supplemented our species-level representative MAG collection “Archaea ESTrep-21” (n=21) with publicly available archaeal MAGs and genomes from the UHGG collections v2.0.2 (n=28)^1^. We performed species-level dereplication for 21 species from the ‘Archaea ESTrep-21’ collection and 28 species from the UHGG collection. Following this process, we compiled a refined set of 37 unique species, which we refer as **‘Archaea GUTrep-37’** (**Figure 2a, Supplementary Table S3**). Of these, 12 species were shared between the “ESTrep-21” MAG collection and the UHGG collection, while 16 species were exclusive to the UHGG collection, and 9 species were unique to the “ESTrep-21” MAG collection (**Figure 2b**).

**Figure 2.**
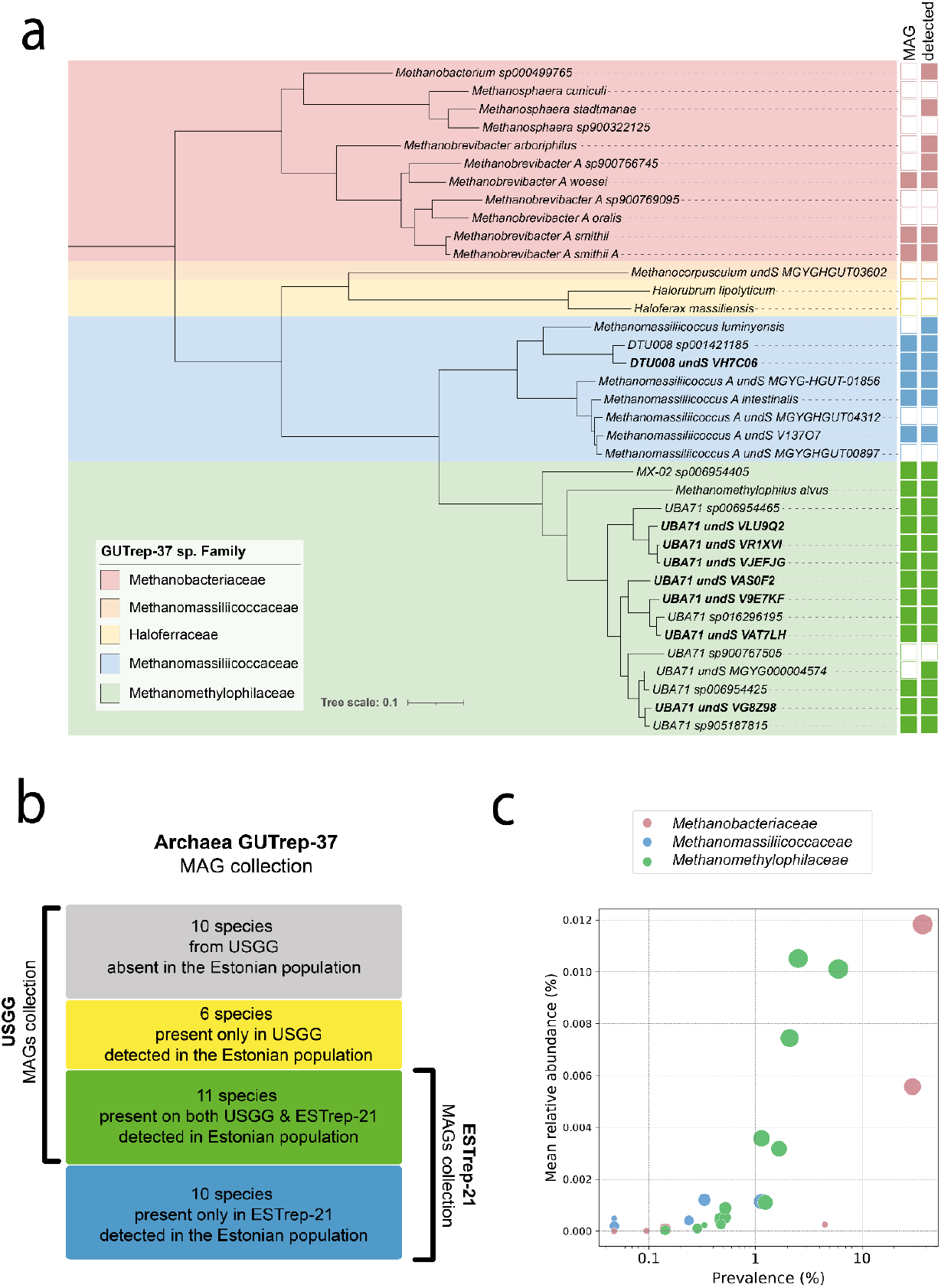
The phylogenetic tree, prevalence and abundance of the human gut archaea in Estonian population. **a.** FastTree approximate maximum-likelihood phylogenetic tree of human gut archaea, including species assembled in this study and those from the USGG (“Archaea GUTrep-37” collection). The tree was constructed using multiple sequence alignments (MSAs) derived from the concatenation of 53 phylogenetically informative markers (arc53) with GTDB-Tk v2.3.0. Phylogenetic tree was constructed with FastTree and generated with iTOL (https://itol.embl.de), with midpoint rooting. Species are labeled with their respective names, while new species are named by their MAG_ID or UDGG_ID. Adjacent to each species label, two indicator boxes provide additional information: “MAG” - the presence or absence of an assembled MAG for the species in the Estonian population (filled and empty boxes, respectively), and “detected” - the detection of the species through mapping in the Estonian population (filled and empty boxes, respectively). Species are color-coded by their family affiliation: *Methanobacteriaceae* (pink), *Methanomassiliicoccaceae* (blue), and *Methanomethylophilaceae* (green), *Methanocorpusculaceae* (orange), *Haloferacaceae* (yellow) **b**. Classification of species from “Archaea GUTrep-37” collection based on their origin and their presence in the Estonian population. **c**. Prevalence and abundance of archaeal species in the Estonian population. Species are color-coded by their family affiliation: *Methanobacteriaceae* (pink), *Methanomassiliicoccaceae* (blue), and *Methanomethylophilaceae* (green).

Read-based community profiling against **‘Archaea GUTrep-37’** archaea MAGs collection showed that 932 out of 1878 faecal samples (49.62%) contained archaea. The mean relative abundance of archaea in the sample was 0.12%, with some samples showing archaea relative abundance up to 3.38% of all reads (**Figure S1**). The prevalence and abundance of archaeal species from the entire Archaea GUTrep-37 collection in the Estonian population are presented in **Figure 2c**. In general, most prevalent and abundant species belonged to the *Methanobacteriaceae* family with 2 species present in more than 10% of samples. *Methanobrevibacter_A smithii* was present in 37% of samples and were shown mean relative abundance 0.01% of all reads and *Methanobrevibacter_A smithi_A* was present in 30% of samples and were shown mean relative abundance 0.006% of all reads. *Methanomethylophilaceae* family also includes species with significant abundance and prevalence. For example, *UBA71 sp006954425* has prevalence 6% and mean relative abundance 0.01%. *Methanomassiliicoccaceae* family is represented by species with lower abundance and prevalence (**Supplementary Table S3**).

Taken together, our population-based metagenomic data help shed light on archaeal diversity, highlighting the Estonian population. Expanded species representative reference **“Archaea GUTrep-37”** may be used in other research for reads based community profiling, especially in northern European populations. “**EstMB MAGdb Archaea-273**” collection is a valuable resource for further archaea research in strain and functional levels.

## Methods

### Description of the samples

In our study, MAGs were recovered samples from Estonian Biobank microbiome cohort, representing Estonian population (EstMB cohort, 2504 samples). For MAGs recovery a subset of samples was used that were sequenced with deep coverage (EstMB deep cohort, n = 1878). All details about samples, DNA extraction and sequencing procedure can be found in a paper by Aasmets et al^9^. In brief, after DNA extraction and DNA library preparation, shotgun metagenomic paired-end sequencing was performed by DNBSEQ-G400RS High-throughput Sequencing Set (FCL PE150) according to the manufacturer’s instructions (MGI Tech, Shenzhen, China). After sequencing human DNA and low-quality sequences were removed.

### MAG assembly and binning

The cleaned reads were assembled into contigs with MEGAHIT v1.2.911^10^. Binning was performed separately for each individual sample. Initially, contigs were binned using single binners: MetaBAT v2.15^11^, MaxBin v2.2.7 (https://sourceforge.net/projects/maxbin2), and VAMB v3.0.7^12^. Given that different binning tools reconstruct genomes differently, a bin aggregation software, i.e., DAS Tool v1.1.4^13^ was employed to integrate the bin predictions from VAMB, MetaBAT2, and MaxBin2. This approach optimized the selection of non-redundant, high-quality bin sets using default parameters. MAGs resulting from this process form the EstMB MAG collection (84,762 MAGs). All MAGs were clustered on species level (ANI index ≥ 95) with dRep^14^ where 69,302 were clustered into 2,257 species clusters and 15,460 MAGs were failed to cluster due to not meeting the built-in dRep quality check threshold.

### Taxonomic annotation

Taxonomy of all representative MAGs and also all MAGs failed to cluster was assigned with GTDB-Tk v2.3.0^15, 16^. Any MAGs that could not be assigned to any known species based on GTDB-Tk v2.3.0^17^ taxonomic annotation were identified as potentially new species.

### Evaluate the quality of MAGs

The quality of all representative MAGs and also all MAGs that failed to cluster was evaluated based on CheckM2^18^ results. Here, high quality (HQ) MAGs exhibited completeness exceeding 90% and contamination below 5%, medium quality (MQ) MAGs failed to classify as a HQ MAGs but displayed completeness greater than 50% and contamination below 10%, while all other MAGs were categorized as low quality (LQ) MAGs.

### “EstMB MAGdb Archaea-273” MAGs collection

For all MAGs classified as archaea (n=316) together with evaluation of completeness and contamination with CheckM218, we determined the genome size and number of contigs with seqkit/2.3.1^19^ and annotated each MAG contigs separately with GUNC^20^. During the subsequent quality control steps, we excluded low quality MAGs (completeness < 50%, contamination > 10%), MAG with abnormal genome size (>2Mb) and MAGs with non-consensus contig annotation (consensus contig annotation refers to the situation where more than half of the contig lengths are annotated with the same genus). This final step helps us eliminate questionable MAGs with unusually large genome sizes and diverse contig origins, even if they may have high completeness and low contamination estimates, leaving 273 archaeal MAGs (Suppl. Table S1).

### “Archaea ESTrep-21” MAGs collection

All MAGs from “EstMB MAGdb Archaea-273” MAGs collection was clustered on species level (ANI index ≥ 95) with dRep^14^, resulting in 21 species level clusters with a representative genome from each cluster. These representative genomes characterise Estonian archaea diversity and form a MAGs collection “Archaea ESTrep-21” which is subset of “EstMB MAGdb Archaea-273” MAGs collection. MAGs from this collection are not stored separately, as they present in bigger “EstMB MAGdb Archaea-273”. If needed MAGs may be sourced from “EstMB MAGdb Archaea-273” collection based on MAG_ID (see Suppl. Table S2). To evaluate strain diversity within “EstMB MAGdb Archaea-273” MAGs collection, MAGs from this collection were also clustered on strain level (ANI index ≥ 99) with dRep^14^.

### “Archaea GUTrep-37” MAGs collection

All MAGs from “Archaea ESTrep-21” MAGs collection was then clustered together with MAGs from publicly available archaeal MAGs and genomes from the UHGG collections (n=28)^1^. Clustering was performed on species level (ANI index = 95) with dRep^14^, for each species cluster representative MAG or genome was selected. As species representative may not be the MAG from Estonian MAG collection and for some species we had only MAGs from USGG collection, we again re-estimate MAGs quality with CheckM2^18^ and MAG/genome characteristics with a seqkit/2.3.1^19^ and store this collection separately to make it available to easy download and use for father mapp based community profiling.

### Phylogenetic analysis

FastTree approximate maximum-likelihood phylogenetic tree was constructed for human gut archaea species, including species assembled in this study and species from the USGG (“Archaea GUTrep-37” collection). Multiple sequence alignments (MSAs) were formed from the concatenation of 53 (arc53) phylogenetically informative markers with GTDB-Tk v2.3.0^15,16^. Visualisation of the phylogenetic tree was prepared with iTOL (https://itol.embl.de), with midpoint rooting.

### Estimation of prevalence and abundance of archaeal species in Estonian population

For mapping-based community profiling we used as a reference “Archaea GUTrep-37” MAGs collection. All reads were mapped against the reference with CoverM (https://github.com/wwood/CoverM) and abundance of all archaea species were estimated for all samples. To estimate species prevalence, we aggregate single sample results to one table (script “parse_abud_results_v2.py”) and perform further calculations with standard python packages.

### Functional annotation

Protein-coding sequences (CDSs), tRNA genes and CRISPR were predicted and annotated with Prokka v.1.14.6^21^ using the parameters ‘--kingdom Archaea’.

### Tools used for data visualization

All visualisations were conducted using python packages pandas, matplotlib, plotly and seaborn.

## Data Records

The MAG sequences from “EstMB MAGdb Archaea-273” MAGs collection have been deposited in the European Nucleotide Archive under study accession PRJEB81541.

## Technical Validation

To improve the quality of our final Archaea MAG collection, we carefully reviewed the initial 316 MAGs. We generated separate figures for the genome sizes of all MAGs, organized by identified orders, and color-coded data points based on contamination levels (**Figure 3**). Notably, we identified 3MAGs that were annotated as unknown species belonging to the order *Korarchaeia*. This looked questionable because, firstly, we did not expect to find archaea from this order in the human gut, and secondly, these MAGs had unusually large genome sizes. In addition, we observed MAGs with anomalously large genome sizes (for archaea from these particular groups) in two other orders, which may also be the result of mis-assembly. Based on these findings, we decided to perform a more detailed manual analysis of all MAGs larger than 2 Mb. We used GUNC^20^ to annotate all contigs in these MAGs and determined a consensus contigs annotation for each MAG. “Consensus contigs annotation” is achieved when more than half of the contig lengths share the same genus annotation. MAGs that did not meet this consensus criterion were excluded from the final set of Archaea MAGs.

**Figure 3.**
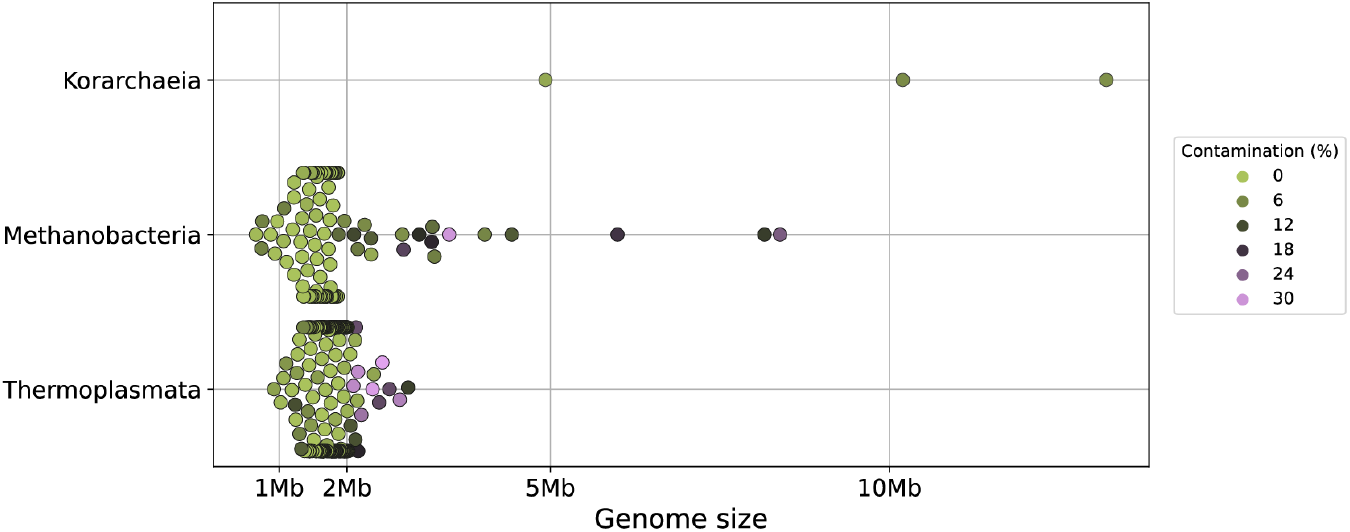
Genome size, taxonomic annotation, and contamination level of initial 316 archaea MAGs. Colour code represents contamination level of each MAG from 0% (light green) to 30% (violet).

## Usage Notes

If dRep is employed as the primary tool for MAG dereplication and clustering, we advise closely examining MAGs that fail to cluster, as our analysis revealed many high-quality archaeal MAGs among these unclustered MAGs. We recommend using the CheckM2^18^ program to assess the completeness and contamination of Metagenome-Assembled Genomes.

## Code Availability

All commands and pipelines used in data processing were executed according to the manual and protocols of the corresponding bioinformatics software. The options and parameters of all tools used for the analysis are described in the main text.

The code for creating figures and tables is accessible on GitHub (https://github.com/Chartiza/ArchaeaDraftGenomes).

## Acknowledgements

We express our gratitude to all individuals who made valuable contributions to the Estonian Microbiome cohort, as well as to those who developed the software and databases utilized in this study. Data analysis was carried out in part in the High-Performance Computing Centre of University of Tartu, and we thank HPC Support Team of the Institute of Computer Science at the University of Tartu for delivering exceptional service and assistance in installing the necessary programs on the cluster. Special thanks to the Writing Retreat organized by the Institute of Genomics, University of Tartu for providing a conducive atmosphere for writing this paper.

This work was funded by Estonian Research Council grant (PRG1414 to E.O) and an EMBO Installation grant (No. 3573 to E.O.)

## Author contributions

K.P. designed the study and pipeline, performed the data analysis, interpreted the data, prepared the figures and tables and wrote the manuscript. E.O. conceptualised and supervised the study, reviewed and edited a draft.

## Supplemental information titles and legends

**Supplementary_Figures_S1. Figures S1**.

**Supplementary Table S1**. “EstMB MAGdb Archaea-273” MAGs collection description. Quality parameters, taxonomic annotation, and genome characteristics (MAG - metagenome assembled genome; HQ-high quality; MQ - medium quality).

**Supplementary Table S2**. “Archaea ESTrep-21” MAGs collection description. Quality parameters, taxonomic annotation and genome characteristics (MAG - metagenome assembled genome; HQ-high quality; MQ - medium quality).

**Supplementary Table S3**. “Archaea GUTrep-37” MAGs collection description. Quality parameters, taxonomic annotation and genome characteristics (MAG - metagenome assembled genome; HQ-high quality; MQ - medium quality).

## References

1. Almeida, A., Nayfach, S., Boland, M., Strozzi, F., Beracochea, M., Shi, Z.J., Pollard, K.S., Sakharova, E., Parks, D.H., Hugenholtz, P., et al. (2021). A unified catalog of 204,938 reference genomes from the human gut microbiome. Nat. Biotechnol. 39, 105–114. 10.1038/s41587-020-0603-3.

2. Chibani, C. M. et al. A catalogue of 1,167 genomes from the human gut archaeome. Nat Microbiol 7, 48–61 (2021).

3. Wampach, L. et al. Colonization and Succession within the Human Gut Microbiome by Archaea, Bacteria, and Microeukaryotes during the First Year of Life. Front. Microbiol. 8, 738 (2017).

4. Volmer JG, McRae H, Morrison M. The evolving role of methanogenic archaea in mammalian microbiomes. Front Microbiol. 2023 Sep 1;14:1268451. doi: 10.3389/fmicb.2023.1268451. PMID: 37727289; PMCID: PMC10506414.

5. Kuehnast, T. et al. Exploring the human archaeome: its relevance for health and disease, and its complex interplay with the human immune system. The FEBS Journal febs.17123 (2024) doi:10.1111/febs.17123.

6. Ghavami, S. B. et al. Alterations of the human gut Methanobrevibacter smithii as a biomarker for inflammatory bowel diseases. Microbial Pathogenesis 117, 285–289 (2018).

7. Mbakwa, C. A. et al. Gut colonization with methanobrevibacter smithii is associated with childhood weight development. Obesity 23, 2508–2516 (2015).

8. Pantiukh, K. et al. Metagenome-assembled genomes of Estonian Microbiome cohort reveal novel species and their links with prevalent diseases. bioRxiv 2024.07.06.602324; doi: 10.1101/2024.07.06.602324

9. Aasmets, O., Krigul, K.L., Lüll, K., Metspalu, A., and Org, E. (2022). Gut metagenome associations with extensive digital health data in a volunteer-based Estonian microbiome cohort. Nat. Commun. 13, 869. 10.1038/s41467-022-28464-9.

10. Li, D., Liu, C.-M., Luo, R., Sadakane, K., and Lam, T.-W. (2015). MEGAHIT: an ultra-fast single-node solution for large and complex metagenomics assembly via succinct de Bruijn graph. Bioinformatics 31, 1674–1676. 10.1093/bioinformatics/btv033.

11. Kang, D.D., Li, F., Kirton, E., Thomas, A., Egan, R., An, H., and Wang, Z. (2019). MetaBAT 2: an adaptive binning algorithm for robust and efficient genome reconstruction from metagenome assemblies. PeerJ 7, e7359. 10.7717/peerj.7359.

12. Nissen, J.N., Johansen, J., Allesøe, R.L., Sønderby, C.K., Armenteros, J.J.A., Grønbech, C.H., Jensen, L.J., Nielsen, H.B., Petersen, T.N., Winther, O., et al. (2021). Improved metagenome binning and assembly using deep variational autoencoders. Nat. Biotechnol. 39, 555–560. 10.1038/s41587-020-00777-4.

13. Sieber, C.M.K., Probst, A.J., Sharrar, A., Thomas, B.C., Hess, M., Tringe, S.G., and Banfield, J.F. (2018). Recovery of genomes from metagenomes via a dereplication, aggregation and scoring strategy. Nat. Microbiol. 3, 836–843. 10.1038/s41564-018-0171-1.

14. Olm, M.R., Brown, C.T., Brooks, B., and Banfield, J.F. (2017). dRep: a tool for fast and accurate genomic comparisons that enables improved genome recovery from metagenomes through de-replication. ISME J. 11, 2864–2868. 10.1038/ismej.2017.126.

15. Chaumeil, P.-A., Mussig, A. J., Hugenholtz, P. & Parks, D. H. GTDB-Tk v2: memory friendly classification with the Genome Taxonomy Database. http://biorxiv.org/lookup/doi/10.1101/2022.07.11.499641 (2022) doi:10.1101/2022.07.11.499641.

16. Parks, D.H., Chuvochina, M., Rinke, C., Mussig, A.J., Chaumeil, P.-A., and Hugenholtz, P. (2021). GTDB: an ongoing census of bacterial and archaeal diversity through a phylogenetically consistent, rank normalized and complete genome-based taxonomy. Nucleic Acids Res. 10.1093/nar/gkab776.

17. Nayfach, S., Shi, Z.J., Seshadri, R., Pollard, K.S., and Kyrpides, N.C. (2019). New insights from uncultivated genomes of the global human gut microbiome. Nature 568, 505–510. 10.1038/s41586-019-1058-x.

18. Chklovski, A., Parks, D. H., Woodcroft, B. J. & Tyson, G. W. CheckM2: a rapid, scalable and accurate tool for assessing microbial genome quality using machine learning. http://biorxiv.org/lookup/doi/10.1101/2022.07.11.499243 (2022) doi:10.1101/2022.07.11.499243.

19. Shen, W., Le, S., Li, Y. & Hu, F. SeqKit: A Cross-Platform and Ultrafast Toolkit for FASTA/Q File Manipulation. PLoS ONE 11, e0163962 (2016).

20. Orakov, A. N. et al. GUNC: detection of chimerism and contamination in prokaryotic genomes. Genome Biology 22, 178–178 (2021).

21. Seemann, T. Prokka: rapid prokaryotic genome annotation. Bioinformatics 30, 2068–2069 (2014).

