## Supplementary Figure S1 for "Human gut archaea collection from Estonian population"

Supplementary figures

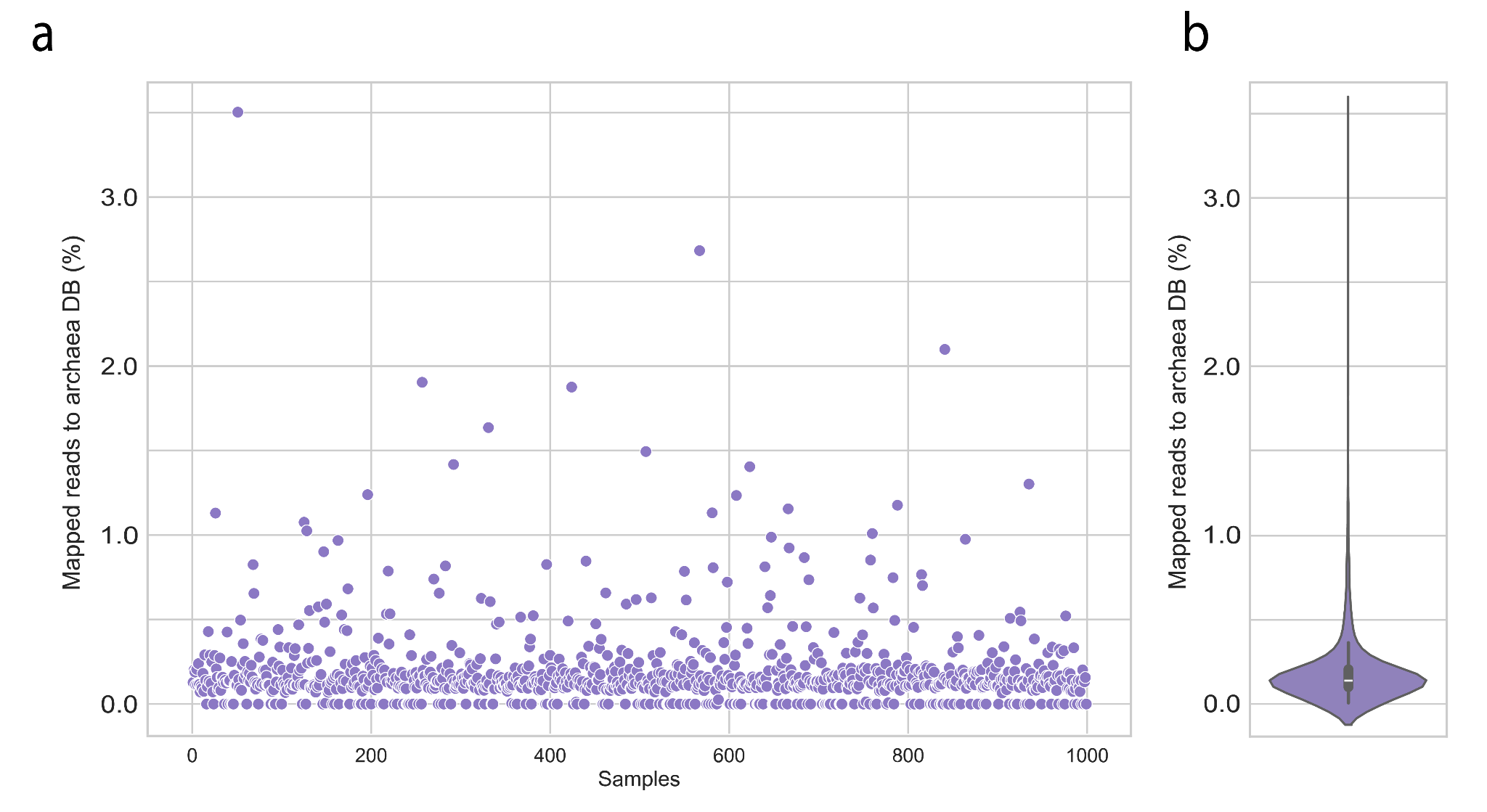

**Figure S1. Relative abundance of archaea in the Estonian population a.** Per-sample relative abundance of all archaea(total number of reads assigned to all archaeal species) **b.** Distribution of the relative abundance of archaea (total number of reads assigned to all archaeal species) in the Estonian population.
